# Attrition Rates in Mindfulness-Based Interventions for Chronic Pain: A Meta-Analysis

**DOI:** 10.1101/2025.05.26.656251

**Authors:** Michael Yufeng Wang, Magelage Prabhavi Perera, Paul B Fitzgerald, Neil W Bailey, Bernadette Mary Fitzgibbon

## Abstract

**Objective:** Mindfulness-based interventions (MBIs) show promise in managing chronic pain but often require substantial time commitments, leading to high attrition rates and concerns about acceptability. This meta-analysis evaluated attrition rates in MBIs for chronic pain and explored factors contributing to participant withdrawal.

**Methods:** Following PRISMA guidelines, 44 studies (45 intervention conditions) were analysed. Data on attrition rates, program characteristics (e.g., delivery method, group vs. individual therapy, session duration), and participant demographics were extracted. Meta-analyses of proportions assessed moderators influencing attrition rates.

**Results:** The overall weighted mean attrition rate was 30.1% (95% CI: 24.5% to 37.3%) with substantial heterogeneity (*I²* = 89.0%). Stricter completion thresholds (defined as the minimum number of sessions required to be considered a programme completer) were associated with higher attrition (*p* < 0.001, *R²* = 28.1%), ranging from an 18% attrition rate for low thresholds (completion of ≥3 sessions required to be defined as a completer) to a 49.7% attrition rate for high thresholds (≥6 sessions). Online delivery showed higher attrition rates (51.0%) compared to in-person delivery (25.6%, *p* = 0.002, *R²* = 17.1%). Individually delivered MBIs were also linked to higher attrition compared to group formats (*β* = 0.216, *p* = 0.039, *R²* = 5.5%).

**Conclusions:** Attrition rates for MBIs in chronic pain management vary widely. Higher attrition is associated with stricter completion criteria, online delivery, and individual formats. These findings highlight the need to optimise MBI programme structure to reduce attrition in MBIs for chronic pain management.

## 1. Introduction

Mindfulness-based interventions (MBIs) are often recommended in the psychological treatment of chronic pain, with empirical evidence supporting their efficacy in reducing pain-related discomfort and functional impairment (Cramer et al., 2012; Greeson & Eisenlohr-Moul, 2014; Scascighini et al., 2008). However, the implementation of MBIs places practical demands on participants, which are often compounded by the challenges of managing chronic pain (Goldsmith et al., 2023; Wilt et al., 2021). Indeed, a significant barrier to undertaking MBIs is the time commitment and sustained effort required to learn and integrate mindfulness practices effectively (Loucks et al., 2022). While the benefits of mindfulness may not be solely dependent on the amount of time spent practicing, MBIs often require participants to engage in sustained and structured mindfulness training to achieve meaningful outcomes, particularly during the early stages of the learning process (Bowles et al., 2022; Davis et al., 2024; Loucks et al., 2022).

The considerable time investment associated with mindfulness training is demonstrated in the program structure of traditional MBIs such as Mindfulness-Based Stress Reduction (MBSR) and Mindfulness-Based Cognitive Therapy (MBCT). Both MBSR and MBCT typically involve weekly 2.5-hour training sessions over eight weeks, alongside daily 45-minute practice sessions for at least six days per week, and these programs often conclude with a full-day retreat (Alsubaie et al., 2017; Parsons et al., 2017). Beyond the initial intensive training period, continued time investment in maintaining a consistent practice regimen is also necessary to help preserve and deepen mindfulness skills (Bowles et al., 2022). Kabat-Zinn (1990), the founder of MBSR, emphasised that mindfulness should be cultivated as a lifelong practice rather than approached as a finite eight-week course. This perspective frames mindfulness training as a long-term lifestyle integration—a skill to be learned, refined, and mastered over time—rather than a single-dose intervention for symptom relief (Milosevic et al., 2020).

Furthermore, the structured design of traditional MBIs, which are taught in a sequence of components that each build on the components taught in the earlier weeks, highlights the importance of program attendance and completion to fully develop the foundational skills of mindfulness (Woods & Rockman, 2021). Partial program completion may not provide sufficient exposure to these core practices, potentially limiting the effectiveness of the intervention. Despite this, program attrition across studies examining MBIs across diverse participant populations and pain conditions varies widely, with rates ranging from 27% to 80% (Brintz et al., 2020; Chen et al., 2023; Ruskin et al., 2017; Simmons et al., 2019). This variability suggests that there may be potential barriers to program completion which could impede participants’ ability to fully engage with and benefit from the MBIs.

Previous research on treatment and clinical trial attrition suggests that variability in program completion is likely influenced by multiple factors, including program design features (e.g., session length and frequency), delivery methods (e.g., online vs. in-person), methodological differences (e.g., inconsistent definitions of completion and attrition rates), and participant-specific factors (e.g., demographic differences or pre-treatment expectations) (van Dijk et al., 2023; Seidler et al., 2021; Skea et al., 2018; Vöhringe et al., 2019). However, for MBIs targeting chronic pain, the specific reasons underlying attrition rates and their variability across studies remain unexplored, as no research has systematically investigated attrition patterns in this context.

Given the important role of program attendance for skill acquisition in MBIs, particularly during the early stages of participation (Loucks et al., 2022), understanding the factors contributing to attrition is fundamental to improving the efficacy of MBIs. This insight can inform refinements to program design and implementation to better retain participants, enable identification of participant characteristics and circumstances that influence program completion, and provide insights into the barriers and facilitators of participation in mindfulness practices in the context of chronic pain management. The present review and meta-analysis therefore aim to systematically evaluate attrition rates in MBIs for chronic pain and explore the moderating factors that influence retention, with the ultimate goal of enhancing therapeutic outcomes and ensuring interventions better meet participant needs and expectations.

## 2. Methods

### 2.1. Protocol and Registration

The protocol for this systematic review and meta-analysis was registered with PROSPERO (CRD42024507477). The systematic review was conducted following the guidelines of the Preferred Reporting Items for Systematic Reviews and Meta-Analysis (PRISMA; Moher et al., 2009).

### 2.2. Search Strategy

The development of the electronic search strategy was conducted in collaboration with the review team. Searches were carried out using the Ovid platform, which included access to databases such as MEDLINE, MEDLINE(R), Epub Ahead of Print, In-Process & Other Non-Indexed Citations, Daily and Versions(R), EBM Reviews - Cochrane Central Register of Controlled Trials, APA PsycInfo, and Ovid Embase.

A set of keywords that aligned with the aims of this review was employed across these databases (e.g., mindfulness, chronic pain). The search terms were tailored to each database to accommodate their unique indexing requirements, with no restrictions on publication date or language. An updated search was conducted prior to the final analysis to include recent studies. A detailed record of the search strategy with a complete list of search terms can be found in supplementary materials.

Two reviewers, MYW and MPP, were provided training prior to conducting the literature screening. This training encompassed the study’s aims, variables, outcomes, and inclusion/exclusion criteria, along with specific search techniques. Training concluded with a pilot review of randomly selected abstracts to ensure inter-rater consistency and reliability in the screening process. The review process was structured in two phases: an initial phase where studies were chosen based on the relevance of their titles and abstracts, and a second phase involving detailed full-text reviews. Any discrepancies in study selection were discussed and resolved with the assistance of two additional independent reviewers, NWB and BMF, to achieve a consensus.

### 2.3. Selection Criteria

The present review sought to examine attrition rates in MBIs for chronic pain management. Studies were included if they met the following a priori criteria: (1) the nature of chronic pain was clearly defined, specifically noting that the condition persisted for more than three months, as per the definition provided by the International Association for the Study of Pain (IASP) (Merskey & Bogduk, 2011); (2) the study implemented MBIs with detailed intervention protocols; (3) the article was published in a peer-reviewed journal and written in English; (4) valid pain measurement tools were included; and (5) participants consisted of an adult population aged 18 years or older. All intervention modalities (e.g., in-person, telehealth, group, or individual) were considered eligible, and no restrictions were placed on the type of pain condition. For studies meeting inclusion criteria but lacking complete attrition data, corresponding authors were contacted to obtain additional information.

Studies were excluded if: (1) they only consisted of purely theoretical content or were commentaries without empirical data, (2) they presented a single case report, (3) they involved participants with prior mindfulness experience, (4) they included interventions that combined mindfulness with other therapeutic approaches (e.g., art therapy, motivational interviewing), (5) they had programs lasting less than two weeks as these were deemed insufficient for evaluating program effectiveness and attrition patterns, (6) they were secondary analysis of previous published studies already included in this review, and (7) they did not present complete data sufficient to enable our analyses, and the corresponding author did not respond to data requests prior to a specified deadline.

### 2.4. Outcome Measures

The primary outcomes of interest covered three key domains: attrition metrics, program characteristics, and participant demographics. Attrition metrics included the number of participants who: (1) initiated the program (defined as attending at least one session), (2) discontinued after attending at least one session, thus not completing the program as designed (e.g., all eight sessions), and (3) attended all sessions of the program. Program characteristics included the mode of the MBI delivery (e.g., in-person, virtual, telephone, hybrid), program duration and frequency, session length, instructor qualifications and experience, total program duration in minutes, and unique protocol characteristics. Additionally, comprehensive demographic information was collected, including participants’ age, gender, race, and pain classification. These variables were selected to provide a thorough understanding of both program implementation factors, as well as participant characteristics that might influence program participation and attrition rates.

### 2.5. Quality of Attrition Reporting

The studies included in this review primarily aimed to assess treatment efficacy rather than feasibility or real-world applicability, which introduces inherent biases in recruitment and attrition considerations. In these studies, attrition is often treated as a factor to minimise rather than an investigable variable. Furthermore, participants tend to be self-selected individuals who express interest in the research, demonstrate willingness to invest time, and show motivation to complete the intervention. These factors may result in both recruitment bias towards participants who are perhaps more able and willing to attend, and potential incomplete attrition reporting across many studies.

Whilst high attrition rates can bias treatment effect estimates and lead to efficacy overestimation (Bell et al., 2013), evaluating the quality of these studies primarily on attrition reporting may not align with their intended objectives. Efficacy studies aim to determine intervention effectiveness under ideal conditions, distinct from effectiveness studies that focus on real-world applicability. This distinction led us to adopt a more balanced approach considering each study’s context and aims. To enhance generalisability, we included all studies meeting eligibility criteria regardless of the quality of attrition reporting. This decision enables a comprehensive analysis of attrition patterns across diverse settings and intervention types. While heterogeneity in attrition rates may partly reflect variations in study protocols—such as session length, which was examined as a potential moderator—this variability may also better represent real-world implementation of psychological interventions.

Rather than conducting a formal risk of bias assessment, we included a dedicated “quality of reporting” discussion section addressing attrition reporting. This section evaluates attrition definition clarity, attendance record completeness, and transparency in the reporting of attrition reasons (if provided). This approach better aligns with the focus of the included studies whilst supporting comprehensive attrition pattern evaluation. The approach also avoids unduly criticising studies based on attrition reporting, since they were not designed to prioritise attrition reporting.

### 2.6. Publication Bias

Publication bias represents a significant threat to the validity of meta-analyses, as studies with statistically significant or larger effect sizes are more likely to be published, while smaller studies with non-significant results are often excluded. This selective reporting can lead to an overestimation of the true effect (Lin & Chu, 2017). To assess potential publication bias, we employed multiple complementary methods: Egger’s regression test to evaluate funnel plot asymmetry, visual inspection of funnel plots to identify distribution anomalies, and the trim-and-fill method to impute missing studies and adjust effect size estimates accordingly.

### 2.7. Statistical Analyses

All statistical analyses were conducted using R version 4.4.2 and the R packages metafor and meta. Given our smaller study pool and observed proportions deviating from 0.5, we implemented the Freeman-Tukey double arcsine transformation (Freeman & Tukey, 1950) to address distribution skewness and achieve more consistent variance. This transformation has demonstrated superior performance compared to alternatives such as logit transformation in simulation studies (Barendregt et al., 2013; Doi & Xu, 2021). Back-transformation was based on Miller’s (1978) equation which facilitated interpretation following significance testing. The analysis employed weighted means using inverse-variance weights, estimated through restricted maximum likelihood (REML) in random-effects models. This approach was justified by heterogeneity indicators from Cochran’s *Q* test (Cochran, 1954) and the *I*² statistic (Higgins et al., 2003; Higgins & Thompson, 2002). The random-effects model was selected for its capacity to account for both within- and between-study variances, which is particularly appropriate given our diverse study methodologies and populations (Wang, 2023).

#### 2.7.1. Sensitivity analysis (outliers)

A sensitivity analysis was conducted to address between-study heterogeneity and evaluate potentially influential outliers. A combination of diagnostic tools and statistical tests were utilised. First, a Baujat plot was used to visually inspect each study’s contribution to heterogeneity (as measured by Cochran’s *Q* test) and its influence on the pooled effect size. This was followed by formal diagnostics, including case deletion methods (e.g., DFFITS and Cook’s distance) and leave-one-out analyses, to evaluate the impact of individual studies on the overall summary effect and heterogeneity. We also screened externally studentised residuals (ESR), using absolute z-scores greater than 2 to identify potential outliers. Rather than automatically excluding identified outliers, we examined the characteristics of studies identified as outliers using the absolute z-scores to understand their influence on the results and potential moderating effects. Outlying studies were retained if they were methodologically sound and were deemed to reflect real-world variability, but excluded if they were deemed to be methodologically flawed or to compromise overall result validity (Wang, 2023). Detailed findings are reported in the results section.

#### 2.7.2 Moderator analysis

A stepwise approach was implemented to investigate moderator effects on attrition rates in mindfulness-based interventions for chronic pain. The analysis proceeded in two phases. Initially, individual meta-regressions were conducted to assess each moderator’s independent contribution to explaining heterogeneity in attrition rates across studies. Subsequently, theoretically related moderators demonstrating statistical significance were integrated into a comprehensive mixed-effects meta-regression model to evaluate their relative contributions while controlling for potential confounding effects. Moderators included demographic variables such as gender, age, and race, as well intervention specific variables such as intervention type, delivery format, program and session length, and trainer qualification. The hierarchical analysis approach enabled a nuanced understanding of both independent and interactive moderator effects on participant retention. For a detailed description and guide of the methods described here please see Wang (2023).

## 3. Results

### 3.1. Study Selection

A total of 4270 studies remained after duplicates were removed. Title and abstract screening resulted in the exclusion of 3,951 studies that did not meet inclusion criteria. Full text screening further excluded 264 studies. There were 26 studies that contained missing or incomplete data, and corresponding authors were contacted via email for supplementary information. Of these, nine authors responded to our request; however, only two of the studies were suitable for inclusion based on our study aims. Finally, 44 studies, comprising 45 conditions, met our inclusion criteria and were included in the subsequent analyses. Figure 1 presents a flow diagram of the study selection process.

**Figure 1.**
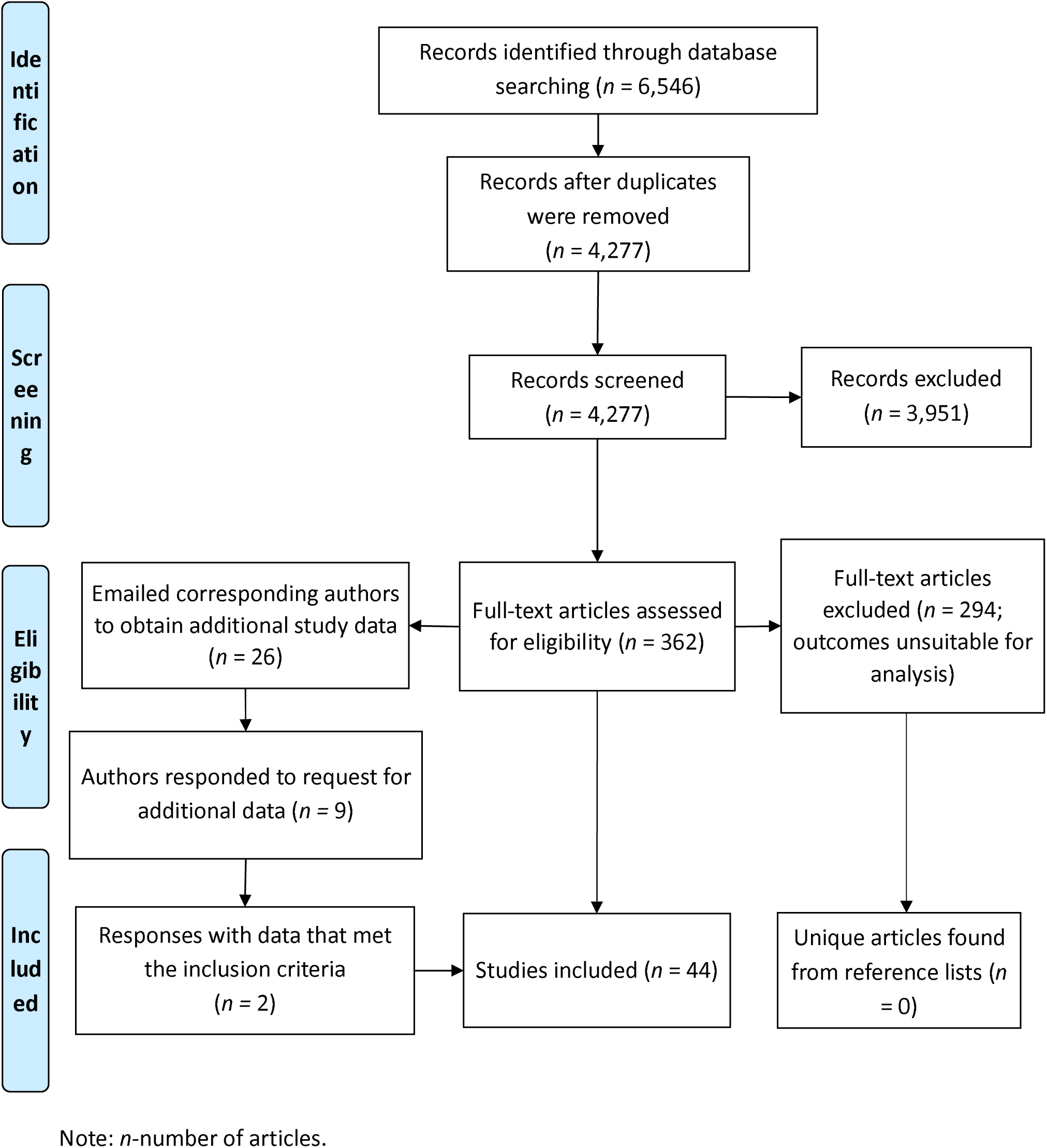
Flow Diagram of Study Selection Process.

### 3.2. Description of Selected Studies

The characteristics of the selected studies are presented in Table 1. As one study featured multiple conditions/arms (comparing MBSR with MBCT) each condition was included as a separate entity to ensure comprehensive evaluation of each intervention approach. Chronic pain conditions across the studies were systematically categorised based on diagnostic similarities, with conditions such as low back pain and back pain consolidated under the broader classification of musculoskeletal pain. The selected studies represented a variety of chronic pain conditions, including musculoskeletal disorders, headaches/migraines, and fibromyalgia, with sixteen studies specifically examining mixed pain conditions.

**Table 1.** Characteristics of Selected Studies.

|  | N of Conditions (%) |
| --- | --- |
| Total | 45 |
| <u>Pain Condition</u> |  |
| Chronic musculoskeletal pain | 14 (31.1%) |
| Fibromyalgia | 8 (17.8%) |
| Headache/Migraine | 5 (11.1%) |
| Mixed - all related to chronic pain | 16 (35.6%) |
| Other | 2 (4.4%) |
| <u>Intervention Type</u> |  |
| MBSR | 28 (62.2%) |
| MBCT | 5 (11.1%) |
| Other | 12 (26.7%) |
| <u>Delivery Mode</u> |  |
| In-person | 37 (82.2%) |
| Telehealth | 2 (4.4%) |
| In-person/Telehealth | 1 (2.2%) |
| Web/Mobile | 5 (11.1%) |
| <u>Therapy Format</u> |  |
| Group | 45 (88.9%) |
| Individual | 5 (11.1%) |
| <u>Program Duration</u> |  |
| 3-4 weeks | 4 (8.9%) |
| 5-6 weeks | 4 (8.9%) |
| 8 weeks | 35 (77.8%) |
| >8 weeks | 2 (4.4%) |
| <u>Trainer Qualification</u> |  |
| Psychologist (clinical, health) | 13 (28.9%) |
| Trained MBSR instructor | 24 (53.3%) |
| N/A | 8 (17.8%) |
| <u>Session Length</u> |  |
| Self-pace online | 4 (8.9%) |
| 1-hr | 2 (4.4%) |
| 1.5-hrs | 7 (15.6%) |
| 2-hrs | 17 (37.8%) |
| 2.5-hrs | 12 (26.7%) |
| 3-hrs | 2 (4.4%) |
| <u>Completion Threshold</u> |  |
| 3-4 sessions | 14 (31.1%) |
| 5-6 sessions | 24 (53.3%) |
| >6 sessions | 7 (15.6%) |
| Homework |  |
| Assigned | 41 (91.1%) |
| Tracked homework completion | 3 (6.67%) |
| <u>Reward</u> |  |
| Yes | 16 (35.6%) |
| No | 29 (64.4%) |

*Note:* MBSR-Mindfulness based stress reduction; MBCT-Mindfulness-based cognitive therapy; MBPM-Mindfulness-based pain management program; Completion Threshold-Program completion criteria used in each study (e.g., number of sessions attended to qualify as a completer); Reward-Whether participants were offered a reward for participation; N of Conditions-Number of unique experimental conditions across all included studies.

Across studies, MBSR was the predominant mindfulness-based approach adopted for chronic pain intervention, typically adhering to standard protocols involving in-person, group-based delivery over an eight-week duration (Santorelli et al., 2017). While some programs included the traditional one-day retreat component, others omitted this element. The most notable variation between MBSR studies appeared in session duration, which ranged from 1.5 to 2.5 hours. Several studies implemented modified versions of MBSR, including brief interventions, extended programs, or adaptations for online or web-based administration.

MBCT was the second most common intervention and followed a similar program structure to MBSR protocols. Other interventions comprised author-developed protocols that followed Kabat-Zinn’s (1994, p.4) definition of mindfulness as “paying attention in a particular way: on purpose, in the present moment, and nonjudgmentally”. Program facilitation was primarily conducted by psychologists and accredited MBSR instructors.

A significant methodological variation was found in the reporting of attrition data across studies. The definition of program completion varied substantially, with some studies classifying participants as completers after attending four sessions (out of the total number of sessions provided by the course which was typically eight) and considering those attending only 1-3 sessions as those have dropped out of the program, with anyone completing 4 or more sessions considered as program completers. Other studies employed more stringent criteria, requiring attendance of at least six sessions or completion of the entire program to qualify as program completers. This heterogeneity in attrition definition was considered a potential moderator (referred to as completion threshold for the remainder of this paper) in explaining variability in attrition rate across studies. Table 1 provides a summary of the characteristics of the selected studies. For a comprehensive description of the selected studies, including detailed methodological specifications, participant characteristics, and intervention protocols, please refer to the supplementary materials.

### 3.3. Overall Attrition Analysis

The meta-analysis, using a random-effects model to account for heterogeneity, revealed a pooled attrition rate of 30.1% (95% CI: 24.5% to 37.3%) across all included studies. Individual study attrition rates ranged from 2.2% to 72.9%. The *Q* statistic was significant (*Q(*44) = 400.510, *p* < 0.0001, *k* = 45), and the *I²* value was 89.0%, indicating significant heterogeneity across the selected studies.

A sensitivity analysis was conducted to identify potential influential studies or outliers. One study was identified as an influential outlier (Pal et al., 2023). However, with this study removed, the pooled attrition rate changed to a negligible extent: 29.6% (95% CI: 23.7% to 35.9%) across the remaining studies. The *Q* statistic remained significant (*Q*(44) = 376.677, *p* < 0.0001, *k* = 44), and the *I²* value was 88.5%. As the methods used in the outlier study were deemed methodologically sound, and the *Q* statistic and *I²* value remained significant and high respectively, the study was retained in further analyses. The persistent significant *Q* statistic and high *I²* value after outlier removal suggest that the observed variability in attrition rates was unlikely to be due to random sampling error alone or influenced by a single outlier. Instead, this variability may reflect differences in population or intervention characteristics, warranting further analysis to examine potential moderators contributing to the considerable variation in attrition rates.

### 3.4. Moderator Analysis

Meta-regression analyses revealed no significant associations (all *p* > .05) between attrition rates and gender, age, race, intervention type, program length, session length trainer qualification, participation reward, or chronic pain condition type (see Table 2 for detailed statistics).

**Table 2.** Meta-Regression Analysis of Moderators on Attrition Rates in Mindfulness-Based Interventions for Chronic Pain.

|  | Q statistic |
| --- | --- |
| <u>Intervention Characteristics</u> |  |
| Completion Threshold | $QM(1) = 13.064, p < 0.001^*$ |
| Pain Condition | $QM(1) = 0.124, p = 0.724$ |
| Intervention Type | $QM(1) = 0.204, p = 0.652$ |
| Delivery Method | $QM(1) = 9.516, p = 0.002^*$ |
| Therapy Format | $QM(1) = 4.271, p < 0.039^*$ |
| Program Duration (in weeks) | $QM(1) = 2.109, p = 0.146$ |
| Trainer Qualification | $QM(1) = 3.627, p = 0.057$ |
| Session Length (in minutes) | $QM(1) = 3.065, p = 0.080$ |
| Total Program Time in Minutes (without homework) | $QM(1) = 1.790, p = 0.182$ |
| Reward | $QM(1) = 1.535, p = 0.215$ |
| <u>Participant Demographics</u> |  |
| % Female (N = 42) | $QM(1) = 0.103, p = 0.748$ |
| M age (N = 42) | $QM(1) = 0.0002, p = 0.989$ |
| % Caucasian (N = 30) | $QM(1) = 0.358, p = 0.550$ |
| *Note:* Completion Threshold- program completion criteria used in each study (e.g., number of sessions attended to qualify as a completer). % of Females-percentage of female participants in the condition; % of Caucasian-participants identifying as Caucasian in the condition; *M*-Mean; Reward-whether participants were offered a reward for participation; Significant Q statistics are highlighted with *. |  |

#### 3.4.1. Completion threshold and attrition analysis

A mixed-effects meta-regression was performed to evaluate the influence of completion threshold on attrition rate. The moderator was found to be statistically significant (*QM(*1) = 13.064, *p* < 0.001). The estimated effect size for the moderator was 0.162 (95% CI: 0.074 to 0.250), explaining 28.1% (*R²* = 28.1%) of the heterogeneity in attrition rates. Figure 2 presents a forest plot of the relationship between completion thresholds and attrition rates. As shown in Figure 2, attrition rates increased progressively with higher completion thresholds. The pooled attrition rate of studies with low completer threshold (participants recognised as completers after completing more than 3-4 sessions) was 18.0% (95% CI: 10.00% to 27.5%), while the rate for the moderate completer threshold (studies that required participants to complete 5-6 sessions) was 31.6% (95% CI: 24.1% to 39.7%). The higher completer threshold, representing participant withdrawal more than 6 sessions, showed the highest attrition at 49.7% (95% CI: 35.2% to 64.3%).

**Figure 2.**
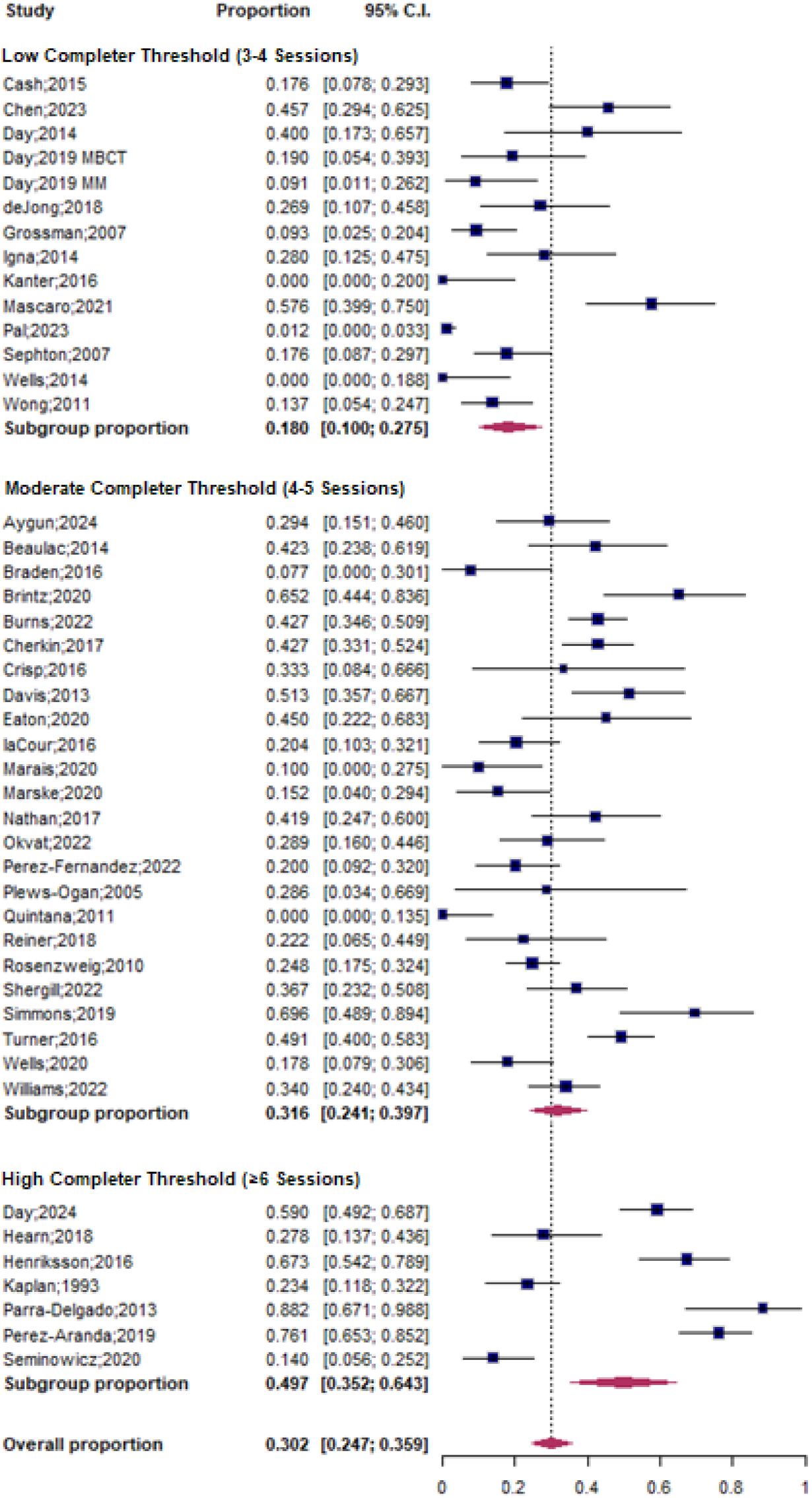
Forest Plot of Attrition Rates by Completion Threshold with Pooled Proportions and 95% Confidence Intervals.

#### 3.4.2. Delivery method and attrition analysis

A mixed-effects meta-regression model was conducted to evaluate the influence of delivery method on attrition rates. Delivery method was found to significantly explain some of the variability in attrition rates across studies (*QM*(1) = 9.516, *p* = 0.002). The in-person delivery method was associated with lower attrition rates (pooled proportion = 25.6%, 95% CI [19.5%, 32.2%]) compared to other delivery methods (e.g., online/mobile or mixed), which had significantly higher attrition rates (pooled proportion = 51.0%, 95% CI [36.2%, 65.6%]). The results indicate that delivery method explains a small but significant portion of the heterogeneity in effect sizes (*R²* = 17.1%). However, as only a small number of included studies (7 studies in total) used the online/mobile or mixed methods, these results should be interpreted with caution. Figure 3 presents a forest plot summarising the pooled attrition rates across different delivery methods.

**Figure 3.**
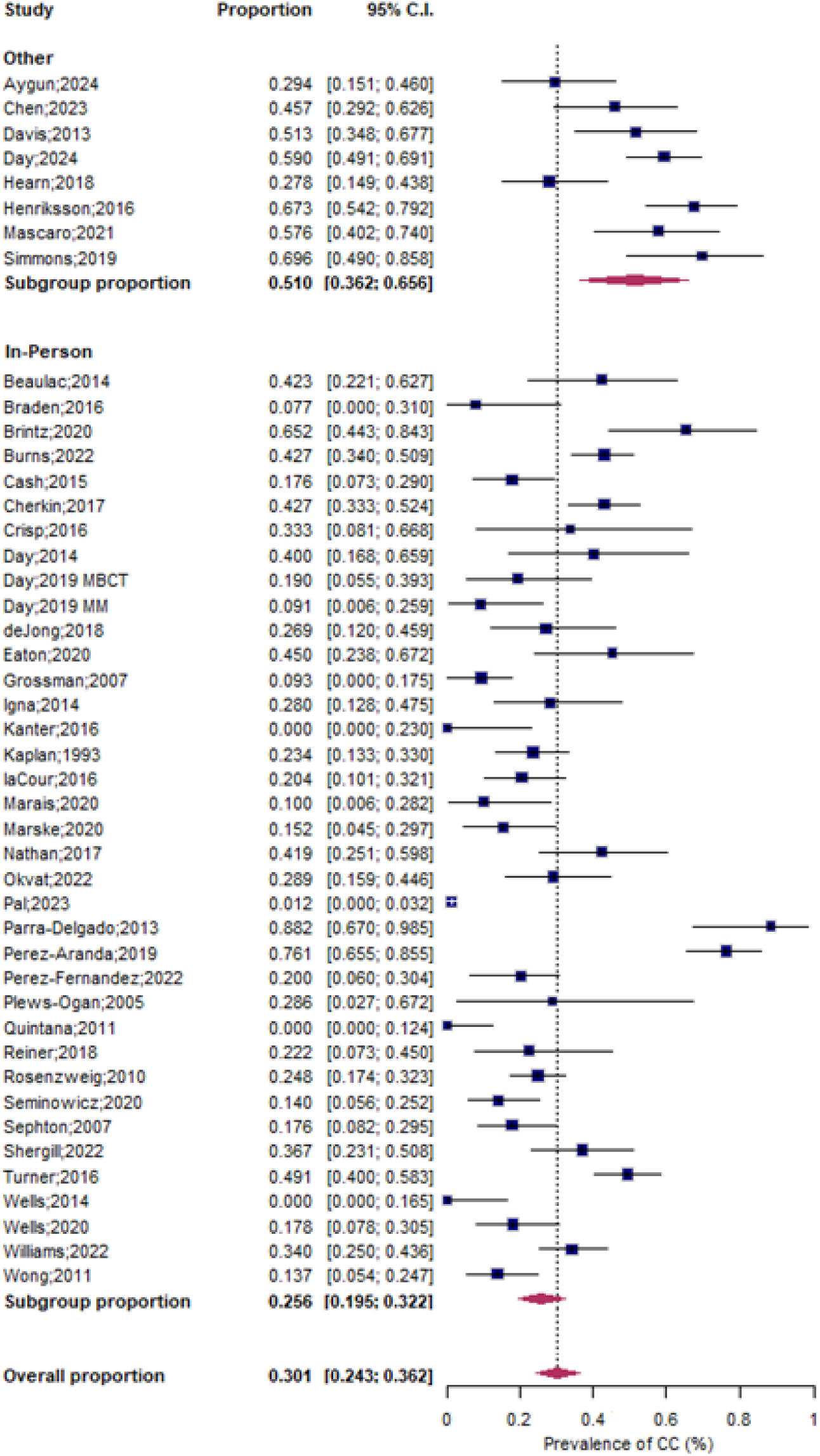
Forest Plot of Attrition Rates by Delivery Method (In-Person vs Other) with Pooled Proportions and 95% Confidence Intervals.

#### 3.4.3. Therapy format and attrition analysis

A mixed-effects meta-regression model was conducted to examine the impact of therapy format (group or individual) on attrition rates across studies. Therapy format was found to significantly explain 5.5% (*R²* = 5.5%) of the variability in attrition rates across studies (*QM*(1) = 4.271, *p* = 0.039). The results indicated that interventions delivered individually, as opposed to group settings, were associated with significantly higher attrition rates (*β* = 0.216, *p* = 0.039, 95% C.I. [0.011, 0.420]). This suggests that group formats may be more effective in retaining participants compared to individual delivery formats. Figure 4 presents a forest plot of attrition rates by therapy format (group vs. individual).

**Figure 4.**
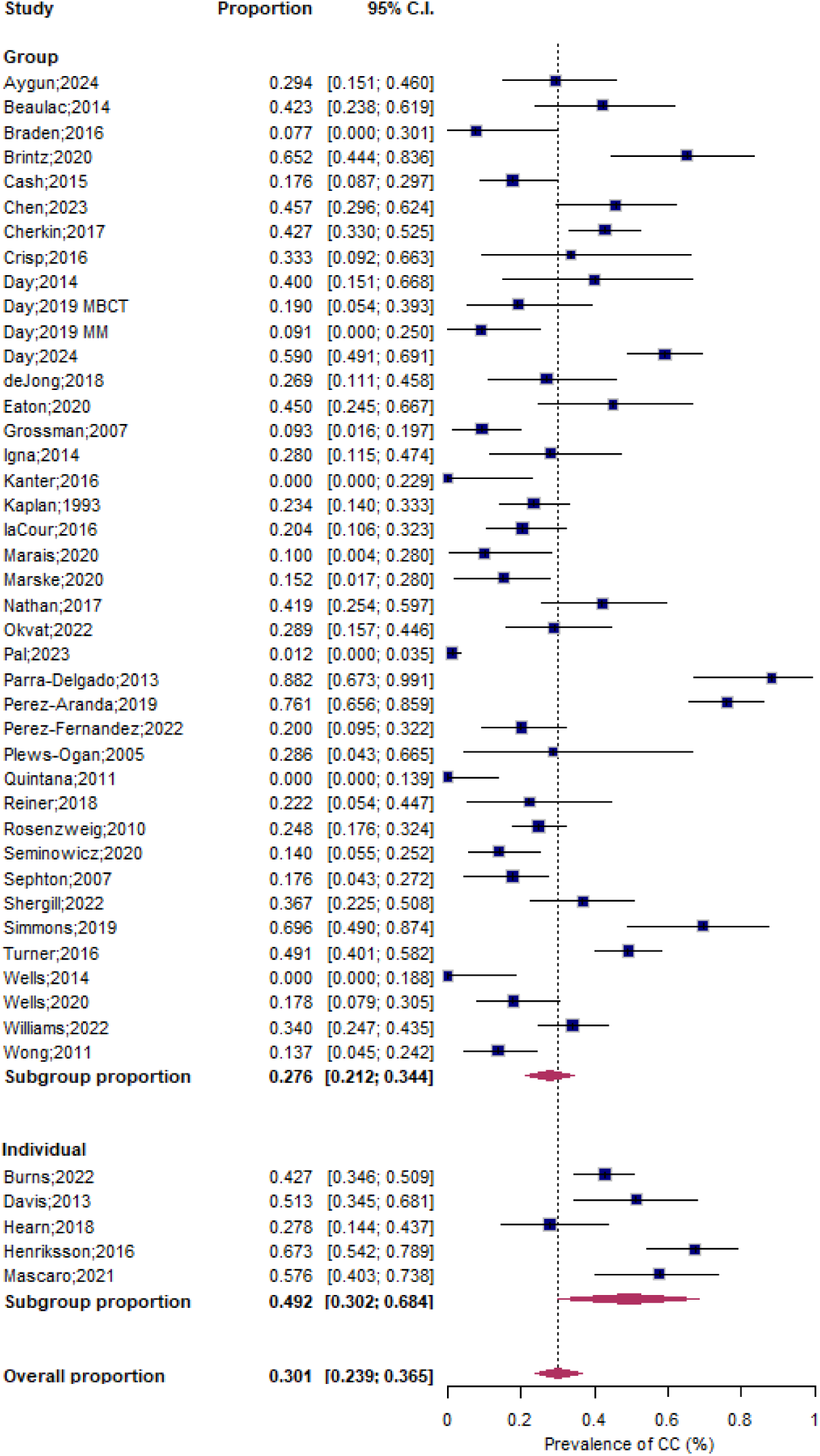
Plot of Attrition Rates by Therapy Format (Group vs. Individual) with Pooled Proportions and 95% Confidence Intervals.

#### 3.4.4. Interaction effects between completion threshold and treatment characteristics

While individual meta-regression analyses identified significant moderators (e.g., completion threshold, delivery method, therapy format), a large proportion of heterogeneity remained unaccounted for. Given the varying definitions of completion thresholds across studies, we conducted a mixed-effects meta-regression model (*k* = 45) to examine potential interactions between completion threshold and other variables. This analysis investigated whether the relationship between completion threshold and attrition rates was moderated by other key variables, particularly delivery method and therapy format, which had shown significant influence on attrition rates.

The mixed-effects meta-regression revealed no significant interactions between completion threshold and either therapy format (*β* = -0.112, *p* = 0.533) or delivery method (*β* = -0.117, *p* = 0.418). This indicates that the relationship between completion threshold and attrition rates remain consistent regardless of delivery method and therapy format. While the main effect of completion threshold remained a significant predictor of attrition rates, substantial residual heterogeneity persisted (*I²* = 84.71%, *Q*(38) = 228.724, *p* < .0001), with the model accounting for 34.2% of the variance (*R²* = 34.2%).

### 3.5. Publication bias

We assessed publication bias using Egger’s regression test, funnel plot asymmetry, and the trim-and-fill method. Egger’s test did not indicate significant asymmetry (*p* = 0.311), while the funnel plot suggested potential bias (figure S1, supplementary materials), with smaller studies showing disproportionate effect sizes. The trim-and-fill analysis estimated 6 potentially missing studies on the right side of the funnel plot, which may reflect an overrepresentation of smaller studies with higher attrition rates. After adjustment, the pooled effect size remained significant (estimate = 0.639, 95% CI: [0.565, 0.712], *p* < 0.0001), indicating that the main conclusions are robust despite potential publication bias. Heterogeneity was high (*I²* = 91.7%, *Q* = 550.375, *p* < 0.0001), suggesting substantial between-study variability, warranting the exploration of potential moderators.

## 4. Discussion

The present study investigated attrition rates in MBIs for chronic pain and identified variables contributing to attrition rates. The general attrition rate of the selected studies (ranging from 2.2% to 72.9%) aligns with findings from efficacy-related meta-analyses, such as that by Marikar Bawa et al. (2015), who reported attrition rates in MBIs for chronic pain ranging from 2% to 50% (median 20%). The meta-analysis conducted by Marikar Bawa et al. (2015) specifically focused on MBSR and MBCT programs lasting a minimum of six weeks. Our meta-analysis included a broader range of mindfulness interventions which provided a wider spectrum of attrition rates. This expanded scope enabled a more comprehensive examination of attrition patterns across diverse intervention types and protocol designs specifically targeting chronic pain. When compared with other meta-analyses across various pain treatment modalities, our findings are broadly consistent. For instance, Oosterhaven et al. (2019) reported attrition rates ranging from 10% to 51% in different pain management programs, while Veehof et al. (2016) found attrition rates between 26.7% and 67.3% across treatments with mindfulness elements—including Acceptance and Commitment Therapy (ACT), MBSR, and MBCT—without finding significant differences between modalities.

Moderator analysis in this study revealed progressively higher attrition rates with increased completion thresholds, particularly in studies requiring participants to have completed 6 to 8 sessions (non-consecutive) to be considered to have completed the program. Variation in delivery method (online versus in-person) and delivery format (individual versus group) explained some heterogeneity in effect sizes, with both online delivery and individual formats showing higher attrition rates compared to in-person and group sessions.

### 4.1. Higher Attrition Rates with Higher Completion Thresholds

An analysis of attrition rates across different completion thresholds revealed that programs requiring attendance at six or more sessions had higher attrition rates. On average, only around 50% of participants attended six or more sessions (non-consecutive). This highlights a significant challenge in retaining participants in MBIs, as half of the participants struggle to meet attendance requirements of programs with higher session counts.

The higher attrition rates observed at higher completion thresholds may stem from several practical and physical challenges, including the severity of disability-related difficulties, logistical constraints, and fatigue, all of which can hinder sustained participation. For example, poor attendance among individuals with chronic pain in MBIs has been linked to difficulties in performing specific mindfulness techniques (Marikar Bawa et al., 2021, 2023). Techniques such as prolonged body scans—which require participants to remain still and focus on various body parts for extended periods—can exacerbate pain-related discomfort and limit the number of sessions participants are able to attend (Marikar Bawa et al., 2021, 2023). Additional factors impacting attendance include individual characteristics such as baseline pain severity, fatigue, and pain in areas that make prolonged sitting difficult, all of which can significantly reduce participation over time, especially in programs with longer session durations (Ellerbroek et al., 2023; Marikar Bawa et al., 2023). However, it is worth noting that no significant differences were found between pain types in our analysis, suggesting that pain types may not directly influence attendance in mindfulness practices. Furthermore, logistical challenges such as transportation difficulties and scheduling conflicts may become more pronounced as programs progress, further increasing the likelihood of missed sessions (Ellerbroek et al., 2023; Wells et al., 2021).

In addition to practical and physical limitations, psychological factors may also play an important role in attrition rates. Research has consistently demonstrated a complex relationship between participant expectations and program retention in MBIs for chronic pain, with unmet expectations of pain relief being a significant contributor to attrition (Day et al., 2016; Marikar Bawa et al., 2021). Participants who enter MBIs expecting mindfulness training to eliminate or significantly reduce pain intensity often discontinue participation when they realise that the program’s focus is instead on pain acceptance and management (Marikar Bawa et al., 2021). To mitigate the impact of these unmet expectations, programs could prioritise clear communication about the objectives of MBIs early in the process. Explicitly explaining, either before enrolment or during the first session, that the program emphasises pain acceptance and coping strategies rather than pain elimination may help align participant expectations with program goals. While this approach may reduce initial enrolment, it could lead to fewer participant withdrawals and better outcomes overall, as participants would be better prepared for the nature of the intervention.

### 4.2. The Role of Session Format and Group Dynamics in Participant Retention

Across the selected studies, programs delivered on an individual basis were associated with higher attrition rates compared to group-based interventions. This finding aligns with evidence from studies examining factors that influence participant experiences in MBIs, which highlight that participants value the group format for fostering a sense of community and shared understanding (Marikar Bawa et al., 2021, 2023). This group setting helped reduce feelings of isolation often experienced by individuals with chronic pain, as mutual validation of experiences creates a supportive environment (Marikar Bawa et al., 2021, 2023). Additionally, group dynamics provide motivational benefits, as peer support and accountability encourage regular attendance and sustained engagement (Ellerbroek et al., 2023). The collective learning environment further enhances participation by allowing individuals to observe and adopt others’ coping strategies, share mindfulness practice adaptations, and engage in collaborative problem-solving (Marikar Bawa et al., 2021). These interconnected mechanisms—social support, shared learning, and collective experience— not only improve participation but also address cost-efficacy concerns, as group formats are generally more cost-effective than individual sessions (Olmstead et al., 2019).

These group-based advantages may also explain the observed differences in retention rates between in-person and online interventions. In general, online programs for pain management have been consistently associated with higher attrition rates compared to in-person programs (Eastwood & Godfrey, 2023; Macea et al., 2010). This disparity may stem from the reduced interpersonal connection in virtual settings, which can diminish the sense of community and engagement. Additionally, online interventions may face challenges such as compromised information delivery, fewer opportunities to clarify program guidelines, and lower accountability (Ball et al., 2020; Ellerbroek et al., 2023). However, it is important to recognise that while group sessions may benefit some participants, the availability of diverse program formats—ranging from intensive in-person retreats to short, accessible sessions on mobile devices—offers flexibility and broadens accessibility. Online MBIs have been shown to improve well-being in certain populations, particularly when tailored to meet individual needs (Alrashdi et al., 2023; Bailey et al., 2018; Macea et al., 2010). Offering a range of delivery methods ensures that MBIs can cater to a wider variety of participant preferences, logistical constraints, and psychological needs, thereby enhancing their accessibility and impact. However, clinicians and researchers should be aware that online delivery methods are associated with higher rates of attrition.

### 4.3. Methodological Limitations and Field-Wide Considerations in Mindfulness-Based Intervention Research

#### 4.3.1. Quality of attrition reporting

While there is considerable focus on establishing the efficacy of MBIs for chronic pain, there is little standardisation in how attrition and retention are defined and reported. The inconsistent definition of completion criteria across studies limits the comparability of studies and leaves us with insufficient insight into the extent to which individuals with chronic pain participate in MBIs. Furthermore, some studies categorise participants as dropouts solely based on failure to complete follow-up outcome assessments, thereby neglecting actual attendance during the intervention. This practice—combined with the overall lack of comprehensive tracking during the intervention (including session-by-session attendance and detailed recording of reasons for discontinuation)—creates a significant gap in our understanding of programme participation and hampers the identification of potential areas for improvement. The absence of reporting standards has broad implications for both clinical practice and research methodology. Without detailed information about when and why participants leave the program, it is challenging to refine programme designs or develop effective strategies to enhance retention.

#### 4.3.2. Lack of homework tracking and methodological limitations

A notable limitation in the selected studies is the lack of systematic tracking and analysis of homework adherence, with only 3 out of the 42 studies reporting on homework adherence and completion rates. This issue is not unique to this meta-analysis but reflects a broader trend across the field, with multiple meta-analyses across the field reporting similar gaps in homework adherence data (Alrashdi et al., 2023; Bermpohl et al., 2023; Konstantinou et al., 2023; Macea et al., 2010). Informal at-home practise is an important component of mindfulness training for chronic pain, as it helps consolidate learning and enhance skill acquisition (Parsons et al., 2017). The lack of homework adherence data undermines our understanding of the dose-response relationship in mindfulness interventions. Without knowing how much participants actually practice, it is difficult to establish evidence-based guidelines for home practice or to identify the barriers that hinder consistent engagement with the recommended practice amounts. This limitation impacts our ability to optimise interventions and develop targeted strategies to support participant with home practice, potentially compromising the overall effectiveness of MBIs for chronic pain management.

Additionally, the attrition rates reported in these studies may not accurately reflect real-world attrition rates due to research-specific confounding factors. For example, the burden imposed by research-related activities—such as pre- and post-intervention interviews and comprehensive questionnaire batteries—may independently influence attrition rates. These components, which are absent in standard clinical implementations of MBIs, make it challenging to isolate intervention-specific factors contribution to program, attrition from those related to research demands. Furthermore, research eligibility may exclude individuals who cannot attend sessions, lack interest, or have conflicting commitments. This pre-screening may limit the generalisability of our findings to broader populations, as it creates a sample that may not fully represent the diversity of individuals who could benefit from MBIs in real-world settings.

Lastly, whilst trainer qualifications were examined as a potential moderator, the analysis was limited to broad categories (psychologist versus trained MBSR instructor) that may not capture important nuances in instructor experience and expertise or the influence of trainer qualifications other than psychologist or trained MBSR instructor on participant retention. However, the effect of trainer qualifications on attrition rates approached our significance threshold (*p* = 0.057), revealing a non-significant trend where programs taught by trained MBSR/MBCT instructors were associated with lower attrition rates. Although the effect of trainer qualifications appears to be small based on these results, a future meta-analysis with a larger sample size may demonstrate a significant effect. Moreover, analyses that incorporate both trainer experience and qualifications may reveal stronger associations, highlighting the importance of considering this factor in future research as a potential determinant of participant attrition rates. A similar near-significant effect was observed for session length, where shorter sessions showed a non-significant trend towards lower attrition rates (*p* = 0.080). Larger meta-analyses again may confirm this association in future. Nevertheless, even if shorter sessions are shown to reduce attrition, researchers and clinicians must carefully balance this finding against the potential reduction in program efficacy associated with shorter session durations.

Several potential factors contributing to heterogeneity in attrition rates between studies couldn’t be analysed due to insufficient data (Bicego et al., 2021). These include participant-related factors such as pain severity, concurrent health conditions, wait-time before commencing the program, work/family commitments, education, and socioeconomic barriers. Future research should systematically document these variables to better understand their impact on participant retention in MBIs for chronic pain.

### 4.4. Future Directions and Conclusion

This study investigated attrition rates and moderators of attrition in MBIs for chronic pain. Our findings identified higher attrition rates reported in studies that required participants to have attended a higher number of sessions before considering them to be program completers. Significant differences in attrition rates were also observed across delivery methods, with online and individually delivered program formats associated with higher attrition compared to in-person group sessions. Future studies should prioritise the standardised reporting of attrition rates and adherence metrics, including session attendance (specifying the number of individuals who attend each session of the program), reasons for discontinuation, and emphasise systematic tracking of homework adherence. This would enhance comparability between studies and provide insights into factors affecting attrition or conversely, retention. Moreover, future studies should integrate with real-world implementation to better understand program feasibility, adherence, and effectiveness across various clinical settings. Together, this work may assist in developing adaptive program structures that accommodate disability-related challenges while maintaining therapeutic integrity. Thereby, enhancing the accessibility and appeal of MBIs for chronic pain management across diverse populations.

